# Whole mouse body histology using standard IgG antibodies

**DOI:** 10.1101/2023.02.17.528921

**Authors:** Hongcheng Mai, Jie Luo, Luciano Hoeher, Rami Al-Maskari, Izabela Horvath, Johannes C. Paetzold, Mihail Todorov, Farida Hellal, Ali Ertürk

## Abstract

Most diseases involve multiple interconnected physiological systems, but histological evaluation of their pathology is currently limited to small tissue samples. Here, we present wildDISCO, a technology that uses cholesterol extraction to enable deep tissue penetration of standard 150 kDa IgG antibodies in chemically fixed whole mice. Combining wildDISCO with whole mouse clearing, we generate whole-body maps of the nervous, immune, and lymphatic systems and show their close interactions throughout the mouse body.

**Graphical Abstract:** 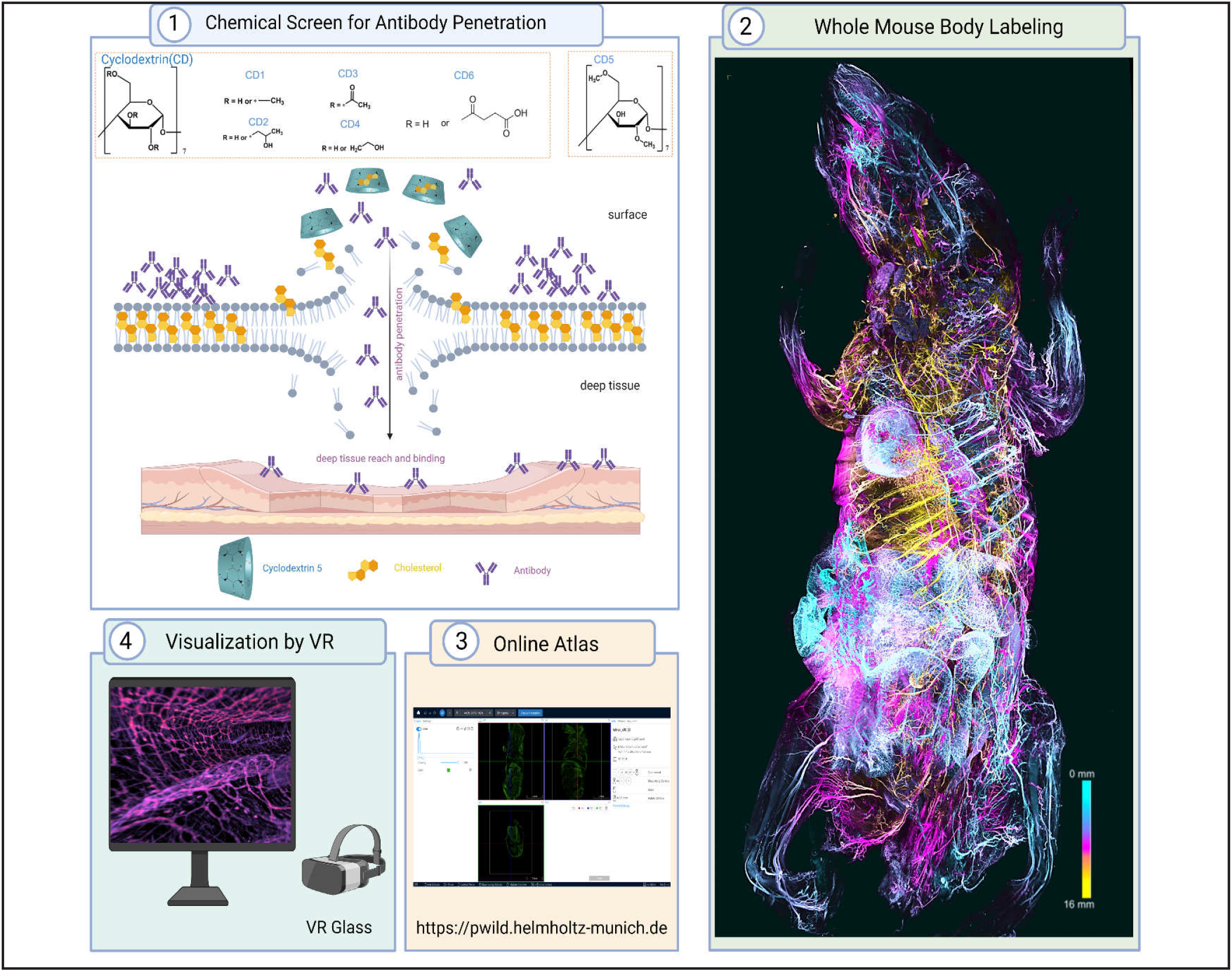

**Highlights:**

1. WildDISCO uses new tissue chemistry based on β-cyclodextrin to enable histology in whole mouse bodies using full-size antibodies
2. WildDISCO generates the first whole mouse body atlases for neurons, immune cells, blood and lymph vessels
3. The whole mouse atlases are available online to study the biological systems in health and disease
4. Virtual Reality (VR) exploration of these atlases disentangles complex anatomical structures between organs and biological systems

**Supplementary Videos can be seen at:** http://discotechnologies.org/wildDISCO/

## INTRODUCTION

More than a century of dedicated work has provided a detailed understanding of the gross anatomy of the human body and the body of common model organisms, and has produced detailed histological maps of many individual organs. However, it remains challenging for a given experimental condition to map the distribution, connectivity and molecular makeup of cell types across the whole body. For example, while the nervous system is connected to every part of the mammalian body, we do not have the cellular level maps of the nervous system to uncover the intricate relationships among organs and between organs and the central nervous system^1-3^. In addition, most methods of imaging nerves or other cells in the context of whole bodies rely on transgenic animals^4, 5^, which severely limits the flexibility of experimental design. Generating new transgenic animals to map changes in the distribution of relevant proteins is usually prohibitively expensive and time consuming. However, such whole-body connectivity maps will be needed to understand the functional interdependence between organ systems and how a disease starting from one part of the body impact the rest such as during neurodegeneration or systemic inflammation.

Recent clearing methods enabled antibody labeling and imaging of intact tissues^6^, mouse organs^7^ and bodies^3, 8-14^, chunk of human organs^15^, and even human embryos^16^, but we still lack suitable, widely-applicable labeling methods for whole mouse bodies. Prior whole-body imaging methods, such as CUBIC, PACT and uDISCO, enabled whole-body imaging, but they relied on transgenic expression of fluorescent proteins in a subset of cells, such as mice expressing Thy-1 EGFP in neurons^17^. vDISCO method5 uses small antibodies called nanobodies (1/10 of IgG size) for whole mouse body labeling. In contrast to the thousands of conventional unconjugated antibodies developed in the last decades, very few nanobodies work in a histological setting. Therefore, a whole-mouse indirect immunolabeling using primary and secondary conventional antibodies would be an invaluable method for many biological applications, including whole-body mapping of cells of interests.

Here, we developed wildDISCO (immunolabeling with wildtype mice and DISCO clearing), a chemical method enhancing the penetration of standard (∼150 kDa size) antibodies into the whole ∼ 2 cm thick mouse body. Our method relies on cholesterol extraction for permeabilization to ensures homogeneous penetration and staining across the tissues of the whole mouse body including muscles, bones, the brain, and the spinal cord. Combining whole-body antibody labeling with DISCO-based tissue clearing allowed us to provide body-wide maps of cell-type and protein distribution with unprecedented ease and will help to advance our understanding of biological systems.

Although homogeneous labeling of whole bodies with small molecules (e.g., DNA-labeling dyes) or nanobodies can be achieved by cardiac pumping of solutions though the mouse vasculature5, this has proven difficult for standard IgG antibodies as 1) the antibodies are degraded and/or precipitate during perfusion, 2) cannot homogenously penetrate different tissue layers including muscles and bones, and 3) the cell membranes are not maximally permeabilized for antibodies to penetrate deep into all tissues with diverse properties.

## RESULTS

### Bone marrow heterogeneity throughout the body

We hypothesized that poor cholesterol extraction from cell membranes might have been the limiting factor for permeabilization. Cyclodextrin is a small oligosaccharide ring use to deplete cholesterol in live membranes^12^. We screened for β-cyclodextrin variants with diverse nature and number of R-motifs (e.g., methyl-, hydroxypropyl-, hydroxyethyl-, succinyl- and acetyl-) **(Fig. 1A)** for their potential ability to facilitate cholesterol extraction in fixed samples in combination with the CHAPS and Triton X-100 detergents to enhance permeabilization **(Fig. S1A)**. Assessing cholesterol extraction using the cholesterol/cholesterol ester-glo assay, we found that the heptakis(2,6-di-O-methyl)-β-cyclodextrin (CD5) extracted most cholesterol from mouse liver tissue after 7 days **(Fig. 1B)**. Addition of CD5 to the permeabilization reactions allowed rapid and homogeneous penetration of methylene blue into the whole mouse brain within 12h, whereas other tested CD chemicals allowed only limited penetration **(Fig. 1C and Fig. S1B)**.

**Figure 1.**
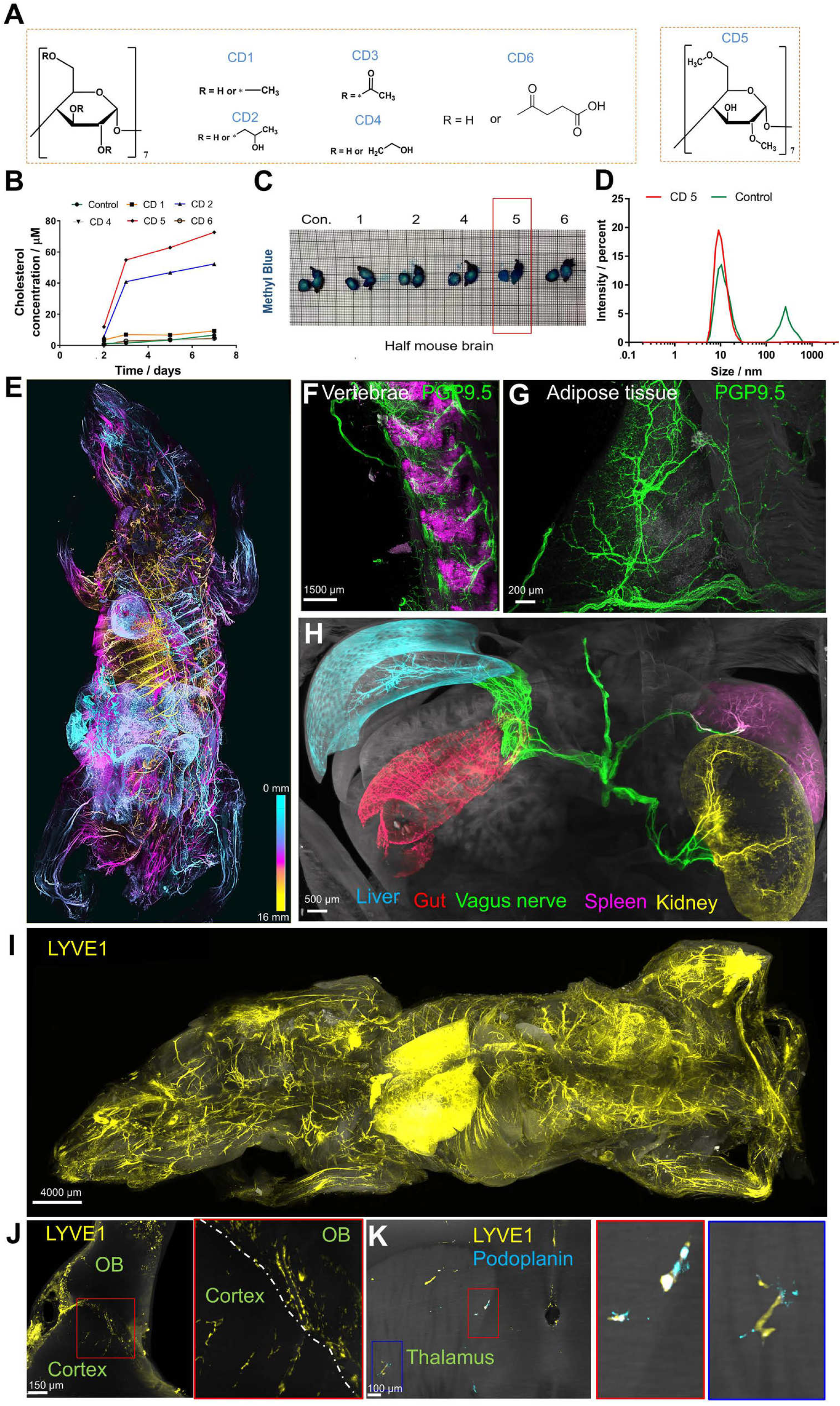
Development of wildDISCO and single antibody whole-mouse staining. (A) The structure of cyclodextrin (CD) with different substituent groups, CD1 (Methyl-β-cyclodextrin), CD2 (2-Hydroxypropyl-β-Cyclodextrin), CD3 (Triacetyl-β-cyclodextrin), CD4 ((2-Hydroxyethyl)-β-cyclodextrin), CD5 (Heptakis(2,6-di-O-methyl)-β-cyclodextrin), CD6 (Succinyl-β-cyclodextrin). (B) Measurements of supernatant cholesterol concentration after different CD-containing buffer incubation at 7th day for 25 mg mouse liver sections. (C) Methylene blue staining of single hemisphere of mouse brains after permeabilization with different CD-containing solutions. CD5 is shown to greatly enhance tissue permeabilization for dye migration compared to others. (D) Dynamic light scattering (DLS) for size distribution of TH antibody in solutions with and without CD5. (E) Deep color coding shows the pan-neuronal marker PGP-9.5+ neuronal projections at different z-levels in the 2.0 cm-thick whole mouse body. (F,G) Details of innervation throughout hard (f, vertebrae) and soft tissues (g, adipose tissue). (H) Manually segmented vagus nerves (green) innervating the liver (cyan), spleen (magenta), intestine (red) and kidney (yellow), highlighted with specific pseudo-colors. (I) A whole mouse stained with a lymphatic vessel marker LYVE1 (yellow). (J) Lymphoid elements (LYVE1) staining was detected in the brain parenchyma of (I) mouse. (K) Mouse brains stained with two different lymphatic vessel markers (LYVE1 and podoplanin) to identify lymphatic endothelial cells were found in the different brain regions.

As cyclodextrins have been previously reported to stabilize proteins in solution by preventing aggregation^18^, we also measured the antibody size in antibody solutions using dynamic light scattering (DLS). We found that after 7 days at room temperature without CD5-containing solution, antibodies showed two peaks in the DLS data, one peak at 11.5 nm presumably for the antibody monomer and a peak for larger sizes, which most likely corresponds to different aggregation states. Addition of CD5 prevented the formation of aggregates **(Fig. 1D)**.

We next tested if the enhanced membrane permeabilization and the decreased aggregation^19^ propensity of antibodies in the CD5-containing buffers increase the homogeneity and depth of antibody staining in whole mouse bodies. Different nerve systems, such as sympathetic and parasympathetic, regulate and coordinate organ function. To reveal delicate nerve innervation of organs in the whole mouse body, we stained the peripheral neuronal network in young adult mouse bodies (∼4 weeks old, ∼10 × 3 × 2 cm dimensions) using protein gene product 9.5 (PGP9.5), a pan-neuronal marker **(Fig. 1E and Fig. 2A and Movie S1)**. After whole mouse antibody labeling using CD5-containing buffers, we rendered them optically transparent and performed panoptic imaging using light-sheet microscopy.

**Figure 2.**
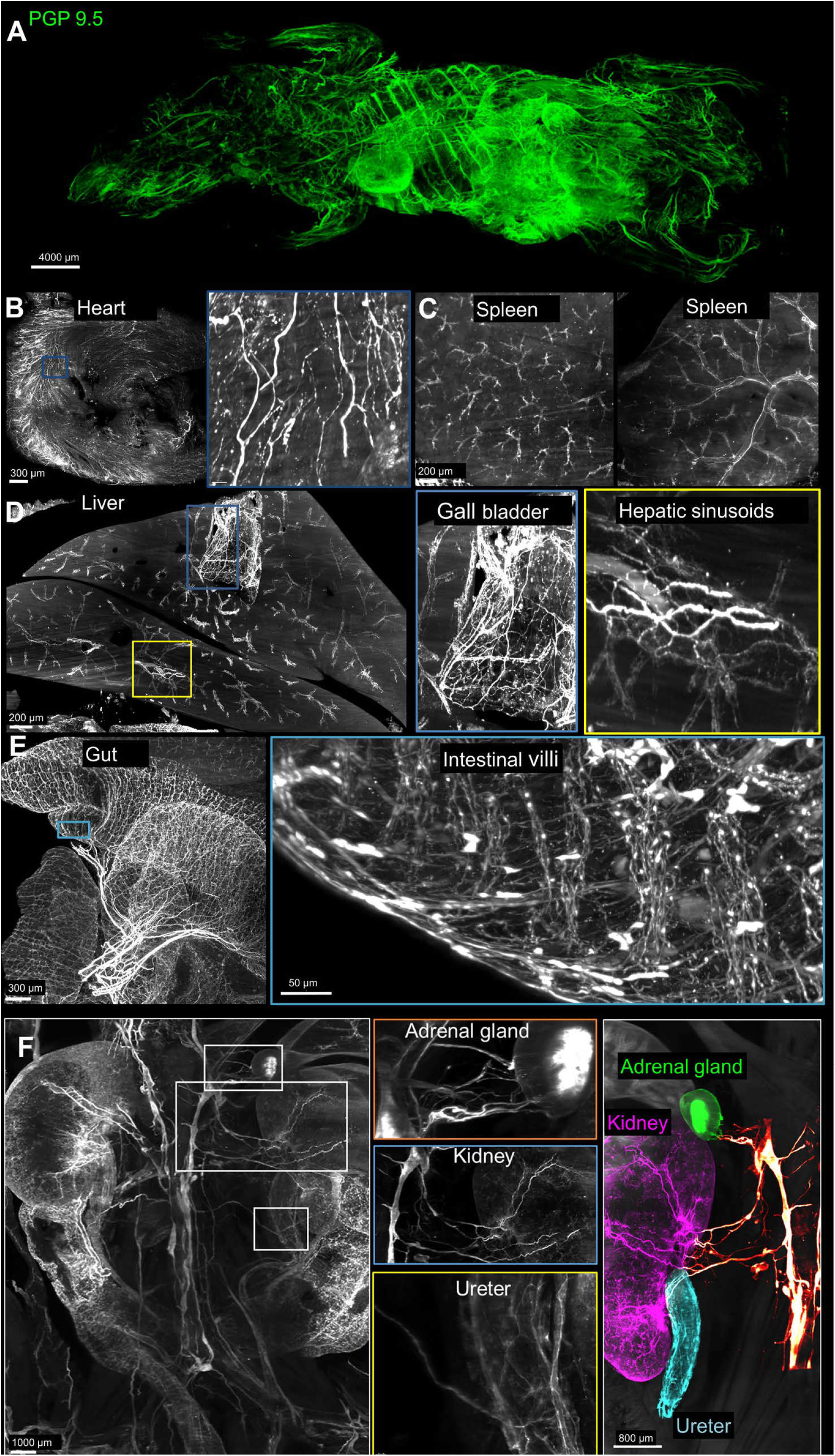
wildDISCO immunostaining of PGP9.5 in the whole mouse body. (A) Maximum projection of peripheral nerve system from a 4-week-old mouse stained with PGP 9.5 antibody using light sheet microscopy. (B-E) Examples of positive PGP 9.5 staining in various organs (heart, spleen, liver, and intestine) with higher magnified areas. (F) Visualization of peripheral nerve innervation on multiple organs (adrenal gland (green), kidney (magenta), and ureter (cyan)).

The peripheral nerve system was homogeneously stained with no systematic differences in signal intensity between tissues as different as vertebrae **(Fig. 1F)** and adipose tissue **(Fig. 1G)** and at different depths of the mouse body.

In the heart, e.g., the network of nerve fibers coursing through the ventricular myocardium was evident **(Movie S2 and Fig. 2B)**. The splenic parenchyma showed nerve fibers’ complex, panicle-like architecture **(Fig. 2C)**. The vagus nerves branched into smaller fiber bundles as it progressed towards the dorsal spleen, where we also visualized the splenic neural network **(Movie S3)**. PGP9.5+ nerve fibers also innervated the hepatic sinusoids **(Fig. 2D)** and distributed along the hepatic duct end-to-end **(Movie S4)**. In the gallbladder, the ganglionated plexus comprising a series of irregularly shaped ganglia **(Fig. 2D and Movie S4)** are clearly visible. In the small intestine, the interconnected ganglionated plexuses on the intestinal wall was observed **(Fig. 2E and Movie S5)**.

In 3D-reconstruction of scans, we could readily observe nerves innervating diverse organs. For example, we could trace vagus nerves, which provides parasympathetic innervation to the abdominal organs, that connects visceral organs, such as the kidneys, adrenal gland, ureter, liver, spleen, and gastrointestinal (GI) tract **(Fig. 1H and Fig. 2F and Movie S6)**, a task that was greatly facilitated with virtual reality visualization techniques. Compared with the whole-organ antibody staining, whole mouse body tracing enabled to visualize nerve connections between different organs **(Fig. 1H)**, which will provide essential clues for understanding the role of nerve communication in normal physiology and disease.

To show the generalizability of the approach, we next stained lymphatic vessels and immune cells using the lymphatic vessel endothelial hyaluronan receptor 1 (LYVE-1) and CD45, respectively.

At the whole-mouse level, we observed the finely structured lymphatic network throughout every part of the body **(Fig. 1I and Movie S7)** and could visualize details of lymph vessel organization in individual organs. For example, LYVE-1+ vessels were observed in the hepatic sinusoidal endothelium and the superficial gastrocnemius **(Fig. 3A and Fig. 3B)**. LYVE-1+ lymph nodes could be observed near the hindlimbs **(Movie S8)**. Especially in adipose tissue, differentially shaped LYVE-1+ cells are seen **(Fig. 3C)**. The larger lymphatic vessels of the kidney branched into lymphatic capillaries with a tree-like architecture **(Fig. 3D and Movie S9)**. Tracheal lymphatic vessels showed a segmental pattern of interconnected vessels **(Fig. 3E)**. In the stomach, lymphatics were unevenly distributed on the gastric walls and had tree-like branches **(Fig. 3F and Movie S10)**. Blunt-ended, tube-like lymphatic capillaries (lacteals) ^20^ were clearly located in the intestinal villi **(Movie S11)**, and the abundant and well-organized lymphatic plexuses and networks were visible on the outer surface of the intestinal wall **(Movie S12 and Fig. 3g-3I)**.

**Figure 3.**
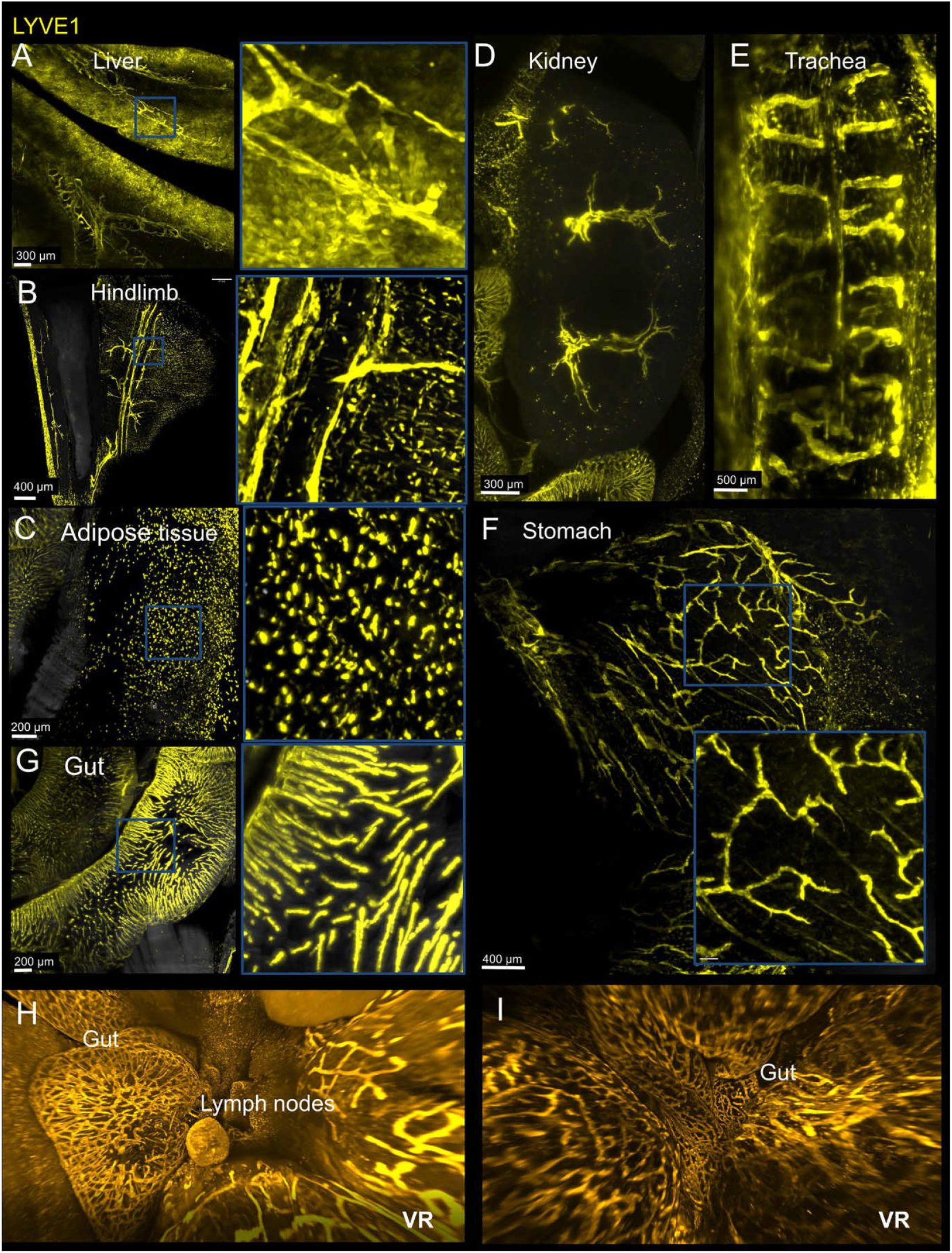
wildDISCO immunostaining of LYVE1 in the whole mouse body. Optical sections examples of whole mouse staining with LYVE1 antibody in different organs, liver (A), hindlimb (B), adipose tissue (C), kidney (D), trachea (E), stomach (F), intestine (G) with higher magnified regions. (H-I) 3D reconstruction view of the intestinal lymphatic network using Syglass reconstruction software.

Previously, the brain parenchyma has been proposed to be devoid of lymphatic vessels^21, 22^, although there is lymphatic drainage from the CNS via meningeal lymphatic vessels. Our whole-body immunolabeling data showed small and short lymphatic capillaries entering the brain parenchyma from the meninges. Some LYVE-1+ lymphatic vessels are also observed to connect the olfactory bulb and the cortex **(Fig. 1J, and Movie S13)**, which was observed by LYVE1 and PROX1 staining, respectively. We also find lymph vessels entering the brain parenchyma around thalamus **(Fig. 1K and Movie S14)**, which was confirmed by both with LYVE1 and Podoplanin staining.

The advantage of wildDISCO compatibility with conventional validated antibodies for labeling allowed us to study the relationship of different physiological systems in the same mouse. First, we co-immunolabeled tyrosine hydroxylase (TH)+ sympathetic nerves and CD45+ immune cells **(Fig. 4A-4C, Fig 5, and Movie S15-S16)**. We find substantial co-localization of immune cells along parts of the vagus nerve, especially at the inferior mesenteric plexus **(Fig. 4A and Fig 4B)**, and frequent contacts between immune cells and sympathetic nerves on the intestinal wall **(Fig. 4C)**.

**Figure 4.**
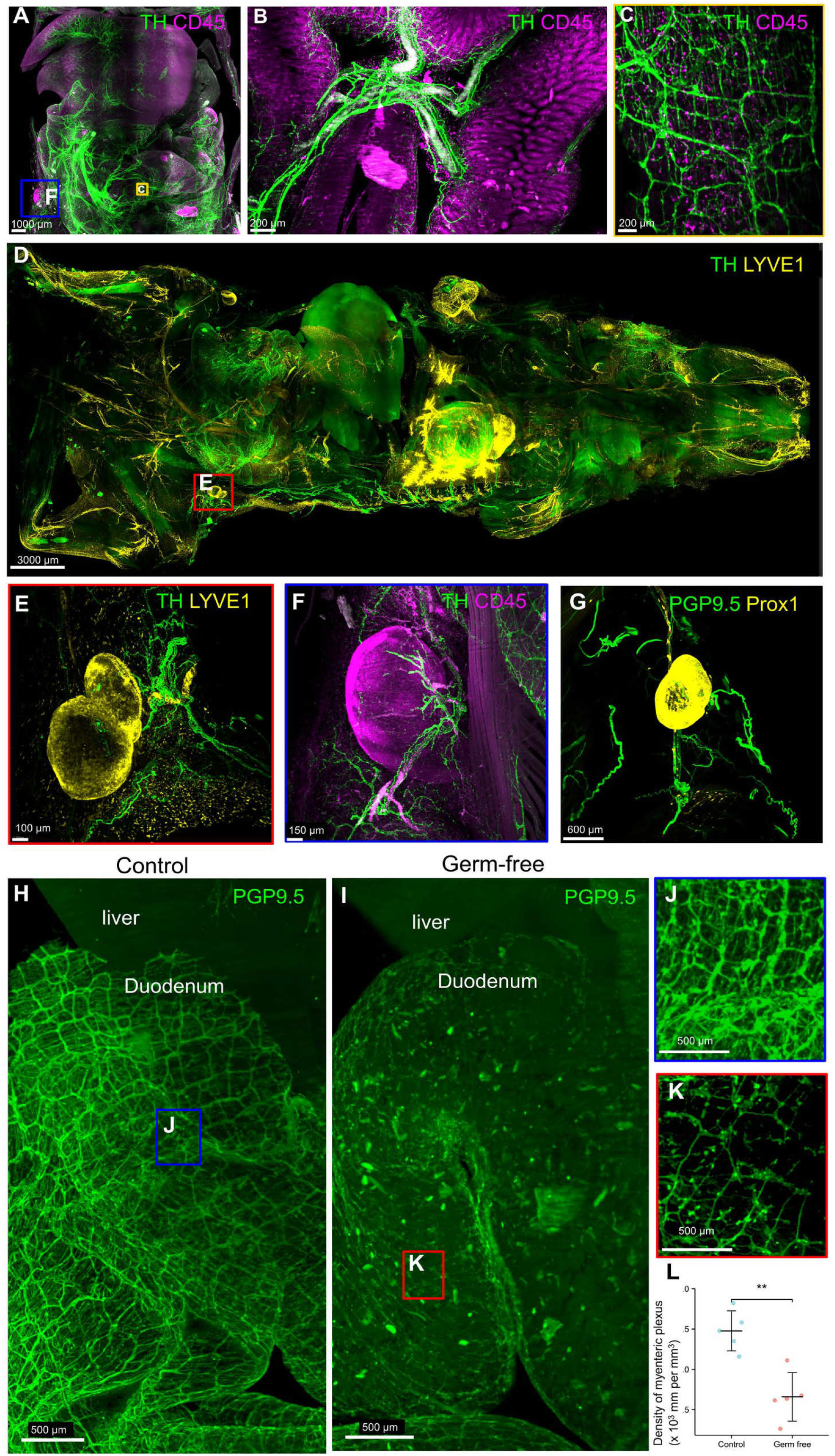
Studying the spatial relationship of different physiological systems using wild-DISCO. (A) Maximum intensity projection of a mouse stained with antibodies against the sympathetic nerve marker tyrosine hydroxylase (TH) (green) and the immune cell marker CD45 (magenta), showing the landscape of neuro-immune interactions in internal organs. (B) The branches of the sympathetic nervous system (TH, green) connect different regions of the intestine. CD45+ cells (magenta) accumulate along parts of the vagus nerve, especially at the inferior mesenteric plexus. (C) High-magnification views of the labeled regions in (A), showing the colocalization of the sympathetic nerve fibers and immune cells on the intestinal wall. (D) Maximum intensity projections of a whole mouse stained with TH (green) and LYVE1 (yellow). (E-G) Representative 2D optical sections of hindlimb LNs stained with TH, PGP 9.5, CD45, Prox1 and LYVE1 as indicated in the images to show the LNs are innervated by peripheral nerves with immunomodulatory potential. (H,I) Representative 3D representation of the myenteric nerve lattice network of WT mice and germ-free mice by immunostaining with antibodies against PGP9.5. (J.K) Higher magnification views of the regions marked by the blue and red boxes in (H) and (J), respectively. In the germ-free mice, the myenteric nerve lattice network appears disorganized, with fewer ganglia. (L) The density of PGP 9.5 myenteric plexus was quantified. n = 5; mean ± SD ; **p< 0.01 (Student’s t test).

**Figure 5.**
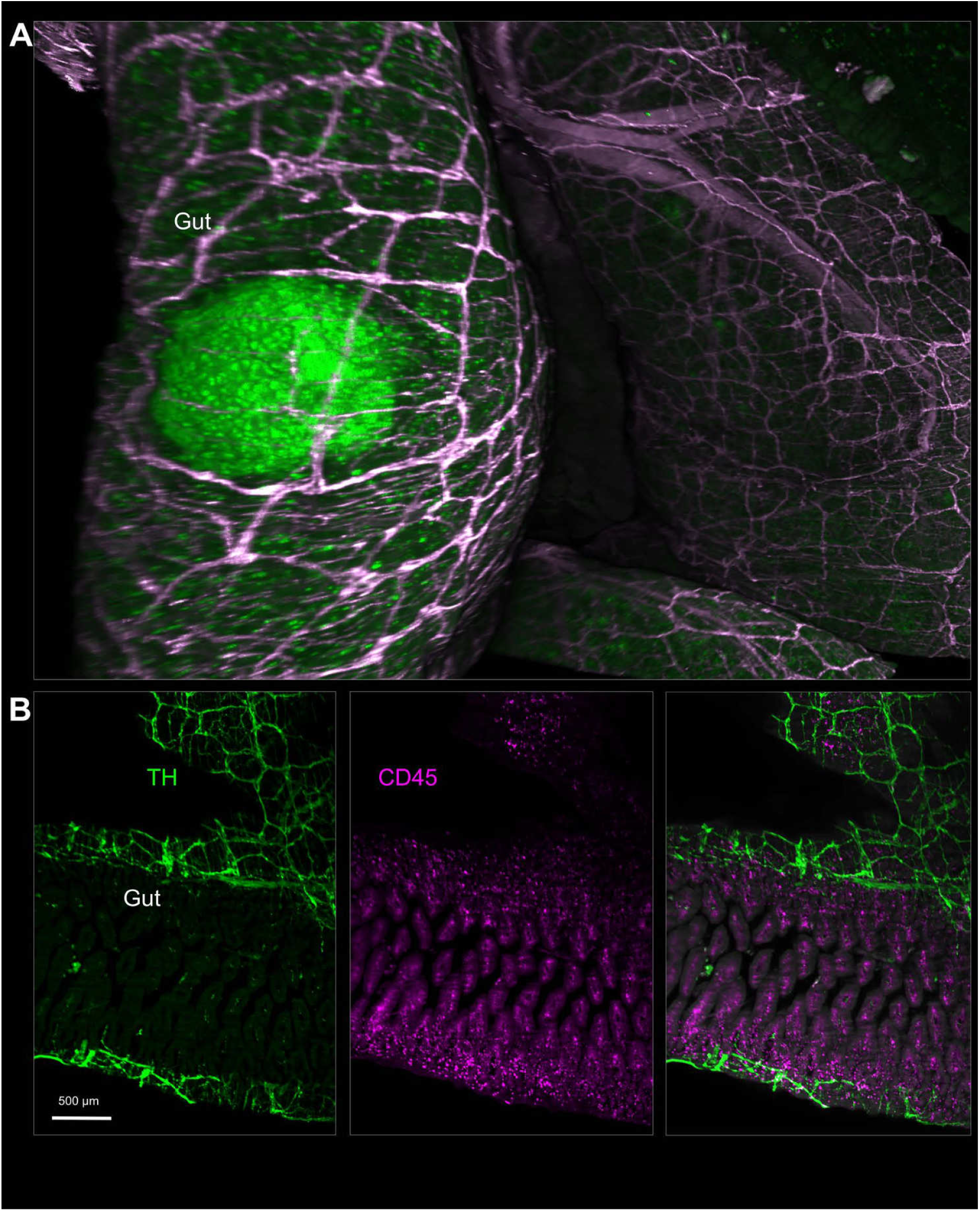
Nerve-immune cell interactions in the gut. (A) 3D reconstruction view of a Peyer’s patch in the intestine stained with CD45 (green) and innervated by TH+ sympathetic nerve (magenta), visualized with Syglass software. (B) Sympathetic nerve marker TH (green) and immune cells marker CD45 (magenta) on the intestinal wall, showed with Imaris software.

To better illustrate neuro-immune interactions in the lymphatic system, especially the lymph nodes (LN), we employed double-staining of nerve fibers and lymphatic vessels **(Fig. 4D, Fig. S2-S3 and Movie S17-S19)**. Among others, large LYVE1+ **(Fig. 4E)** and PROX1+ (Prospero homeobox protein 1, a marker for lymphatic endothelium) **(Fig. 4G)** LN were detected in the mouse hindlimbs **(Fig. 4E, Fig. 4G, and Movie S20-S21)**. CD45 staining confirmed that the observed structures are LNs **(Fig. 4F)**. Co-staining with the pan-neuronal marker PGP9.5 or the peripheral sympathetic neuronal marker TH showed neuronal processes innervating the LN.

Next, we used wildDISCO to assess the effects of biological perturbations. To this end, we compared the structure of the gut-associated nerve system of germ-free mice to specific-pathogen-free (SPF) standard mice. Our double staining data of nerve with lymphatics and nerve with immune cells already showed the enteric nervous system gut intricate details throughout the gut **(Fig. 4A-4D)**. Studing germ-free mice, we found that the PGP9.5+ nerve lattice network is substantially less dense compared to wildtype mice. The density of myenteric plexus reduced from 1.478 (×10^3^ mm per mm^3^) to 0.659 (×10^3^ mm per mm^3^) **(Fig. 4H-4L, Fig. 6 and Movie S22)** confirming the importance of gut microbiota interaction for development and /or maintenance of the mesenteric plexus^23^.

**Figure 6.**
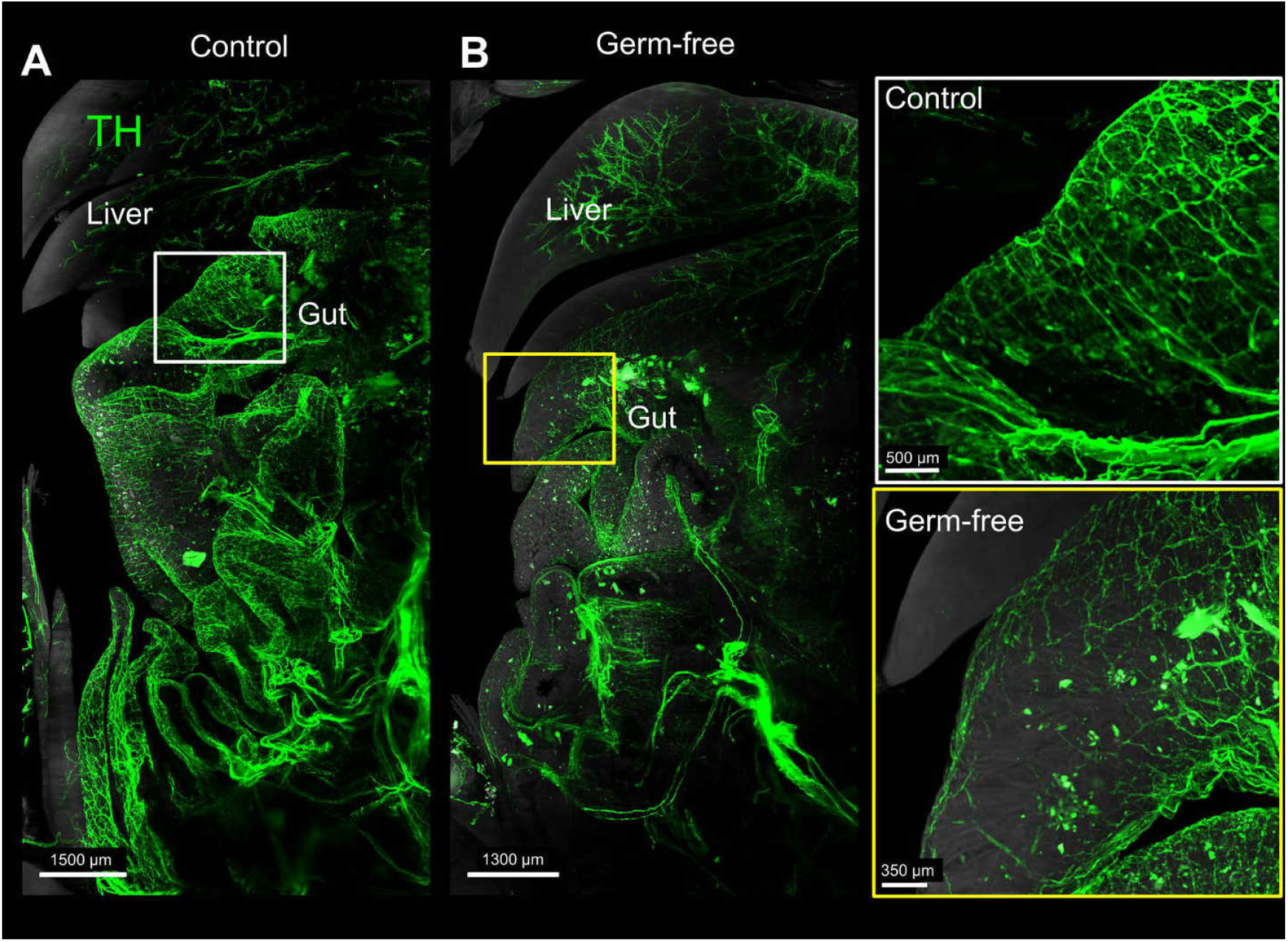
Influence of the microbiota on sympathetic nerve of the mice. (A-B) The myenteric nerve lattice network of WT mice and germ-free mice by immunostaining with antibodies against TH. Higher magnification views of the regions marked by the white and yellow boxes, respectively.

## DISCUSSION

Here we show that unbiased imaging of transparent whole mouse bodies at cellular resolution provides a comprehensive view of biological systems (nerve or lymphatic systems) in health and disease. WildDISCO does not rely on the transgenic expression of fluorescent proteins as it permits the use of off-the-shelf IgG antibodies to homogenously and simultaneously staining structures in the whole mouse body. The mouse head is a perfect example of the versatility of the method as it combines hard (skull) and soft tissues (brain). Using wildDISCO, we could map all the lymphatic vessels in and around the brain parancyma in intact mouse heads.

In summary, wildDISCO technology achieves a homogeneous and simultaneous antibody staining throughout the entire mouse bodies (a summary of results shown in **Movie S23**). Aided by the VR visualization, previously inaccessible 3D anatomical information becomes possible, allowing a more comprehensive understanding of the initiation, progression, and extent of pathologies at the whole organism level in mice (example VR movies are **Movies S24-33**).

## MATERIALS AND METHODS

### Animals involved in the study

We used the following mix-gender animals for the wildDISCO study: 4-week-old wildtype mice (C57BL/6J, CD1 and Balb/c) purchased from Charles River Laboratories. Animals were housed on a 12/12 hr light/ dark cycle and had random access to food and water. Temperature was maintained at 18-23 °C and humidity was at 40-60%. Age- and sex-matched C57BL/6J germ-free mice were purchased from the Technical University of Munich (Institute of Nutrition and Health, Core Facility Gnotobiology), and were housed in a germ-free isolator house. The absence of bacteria was confirmed in the germ-free mice by microbial cultures, and mice were then used for further experiments. Each antibody was repeated successfully on at least five mice and also by at least three different person. Animal experiments were performed according to the institutional guidelines of the Ludwig Maximilian University of Munich and the Helmholtz Munich Center German Mouse Clinic after approval of the Ethical Review Board of the Government of Upper Bavaria (Regierung von Oberbayern, Munich, Germany).

### Screening of cyclodextrin-containing buffer for cholesterol extraction

Cholesterol extraction was measured using the cholesterol/ cholesterol Ester-GloTM assay (Promega, Madison, USA). 25 mg of PFA fixed mouse liver was incubated in 3 ml of 1% w/v different cyclodextrin-containing antibody buffers: 2-Hydroxypropyl-β-Cyclodextrin (PanReac AppliChem, A0367,0100), Methyl-β-cyclodextrin (Sigma-Aldrich, 332615-25G), (2-Hydroxyethyl)-β-cyclodextrin (Sigma-Aldrich, 389137-10G), Triacetyl-β-cyclodextrin (Sigma-Aldrich, 332623-10G), Succinyl-β-cyclodextrin (Sigma-Aldrich, 85990-500MG) and Heptakis(2,6-di-O-methyl)-β-cyclodextrin (Sigma-Aldrich, 39915-1G). The assays were measured at different time points (2d, 3d, 5d, and 7d). 5 ul aliquot of the supernatant was diluted 10-fold in cholesterol lysis solution and incubated for 30 min at 37 °C. Cholesterol detection reagent was then added to the samples and incubated for 60 minutes at room temperature. The value was measured using a Centro LB 96 plate reading luminometer (Berthold, Bad Wildbad, Germany).

### Evaluation of different cyclodextrin-containing buffer effect on the stabilization of antibody

The homogeneity of antibody, in other words of antibody aggregations, in different cyclodextrin-containing buffers was measured by dynamic light scattering (DLS). TH primary antibody was selected to evaluate antibody stabilization and homogeneity. TH antibody (Millipore, AB152) (Mw: 150kDa, Concentration: 10g/l) was dissolved in buffer with and without Heptakis(2,6-di-O-methyl)-β-cyclodextrin (Sigma-Aldrich, 39915-1G) (1% w/v) at room temperature. After 7 days of incubation, the buffer solutions were diluted and afterwards measured in a folded capillary cell (DTS 1070) using a Zetasizer Nano ZS (Malvern, Worcestershire, UK). Samples were measured three times with six sub runs each. The temperature was set to 25 °C.

### Permeabilization capacity of different cyclodextrin-containing buffer on half brain using methylene blue

Mouse half brains were incubated with various cyclodextrin buffers at 1% (w/v), 45mL for 3 days at 37°C. After PBS wash twice, the samples were added 45 mL of 0.03% methylene blue and incubated overnight at 37°C.

To determine the efficiency of methylene blue staining after incubation with different CD buffers, samples were cut in half in the middle line to evaluate the efficacy of the inner tissue staining. The camera images of the samples were analyzed by ImageJ for profile plot along and the pixels were quantified under threshold gray value.

### Perfusion and whole mouse body fixation

Mice were deeply anesthetized with (0.05 mg/kg fentanyl, 0.5 mg/kg medetomidine, and 5 mg/kg midazolam with intraperitoneal injection) and perfused intracardially with heparinized 0.01 M PBS (10-25 U/ml final heparin concentration, Ratiopharm, N68542.03; perfusion volume 12 ml/minute with an ISMATEC peristaltic pump system). After washing out the blood of the mice for 5-10 minutes, 4% paraformaldehyde (PFA) in 0.01 M PBS (Morphisto, 11762.01000) was perfused 10-20 minutes. The mouse bodies were skinned and transferred to 0.01 M PBS after post fixation in 4% PFA for 6 hours at 4°C.

### wildDISCO whole-body immunostaining, PI labeling and tissue clearing

The wildDISCO whole-body immunostaining protocol is mainly based on a setup for pumping the pretreatment solutions and immunostaining buffers through the mouse heart and vasculature to perfuse the whole body. The pumping setup has been previously described6,13. In brief, after post fixation of PFA and 0.1 M PBS washing twice for 30 mins, the mouse body was placed in a 300ml glass chamber and the perfusion needle was inserted into the mouse heart through the same hole as in PFA perfusion. Then, the perfusion needle was connected to an ISMATEC peristaltic pump (REGLO Digital MS -4/8 ISM 834; reference tube, SC0266), which maintained pressure at 160-230 mmHg (45-60 rpm) and was used to establish transcardiac circulation. The pump was equipped with two channels. One was used to pump the solution through the heart to circulate throughout the mouse, while the second channel collected and circulated the solution leaving the mouse body. In the first channel, a 1-ml syringe tip (Braun, 9166017V) was used to connect the perfusion needle (Leica, 39471024) and the reference tube (Ismatec Reglo, SC0266) which is from the pump and set for circulation of the solution through the heart into the vasculature. Since the second channel allowed the solutions to recirculate, the inflow tubing was immersed in the solution chamber of the glass chamber. After the pump and channels are set up, the needle tip was fixed with superglue (Pattex, PSK1C) to ensure continuing and stable perfusion. All of the following perfusion steps were performed using the setup explained above. The mice were first perfused with 0.1 M PBS overnight at room temperature, followed by a 2 day perfusion with the decalcification solution containing 10 w/v% EDTA (Carl Roth, 1702922685) in 0.1 M PBS, and the pH was adjusted to 8-9 with sodium hydroxide (Sigma-Aldrich, 71687) to decalcify all bones at room temperature. Then, the mouse body were perfused three times with 0.1 M PBS and washed for 3 hours each time. Next, the mouse was perfused for 1 day with the permeabilization and blocking solution containing 10% goat serum and 2% Triton X-100 in 0.1 M PBS. Then the mice body were perfused with the primary antibodies TH (Millipore, AB152), PGP9.5 (Proteintech, 14730-1-AP), LYVE1 (Thermo Fisher Scientific, 14-0443-82), CD45 (BD Biosciences, 550539), PROX1 (Abcam, ab101851), Podoplanin (Abcam, ab109059) (25 µg in 250 ml, diluted 1:10,000) or 290 µl PI (stock concentration 1 mg ml-1) incubated for 7 days with the 250 ml immunostaining buffer containing 3% goat serum, 10% CHAPS, 2% Triton X-100, 10% DMSO, 1%glycine, 1% Heptakis(2,6-di-O-methyl)-beta-cyclodextrin in 0.1 M PBS. The mouse body was then washed three times in 0.1 M PBS and each time proceeded 12 hours at room temperature. Then, the mice bodies were perfused in the immunostaining buffer at room temperature with the Alexa fluorescent dye-conjugated secondary antibodies: Alexa Fluor 647 goat anti-rabbit IgG antibody (Thermo Fisher Scientific, A-21245) or Alexa Fluor 647 goat anti-rat IgG antibody (Thermo Fisher Scientific, A-21247) (25 µg in 250 ml, diluted 1:10,000) for 7 days. The mice bodies were washed three times with 0.1 M PBS, each time 12 hours. After the immunostaining steps are done, the mice were transferred to a fume hood and were cleared using the 3DISCO passive whole-body clearing protocol as previous reported9. Briefly, mice bodies were placed in a 300 ml glass chamber and immersed in 200 ml of the following gradient of THF (Tetrahydrofuran, Roth, CP82.1) in distilled water with gentle shaking: (50% x1, 70% x1, 80% x1, 100% x2, 12 h for each step), followed by 3 h in dichloromethane (DCM, Sigma, 270997) and finally in BABB solution (benzyl alcohol + benzyl benzoate 1:2, Sigma, 24122 and W213802) until the bodies were optical transparent.

### Light-sheet microscopy imaging

Image stacks were acquired using a Blaze ultramicroscope (LaVision BioTec GmbH, version 7.3.2) with an axial resolution of 4 μm and the following filter sets: ex 470/40 nm, em 535/50 nm; ex 545/25 nm, em 605/70 nm; ex 640/40 nm, em 690/50 nm. Whole mouse bodies were scanned individually with an Ultramicroscope Blaze light sheet microscopy 4x objective (Olympus XLFLUOR 4x corrected/0.28 NA [WD = 10 mm]). We covered the entire mouse in with 9×23 tile scans with 20% overlap and imaged separated from the ventral and dorsal surfaces to a depth of 10 mm, covering the entire body volume with a Z-step of 10 µm. The width of the light-sheet was reduced to 60% to achieve maximum illumination of the field of view, and the exposure time was set to 120 ms. The laser power was adjusted as a function of the intensity of the fluorescence signal to avoid saturation. The acquired raw images TIFF were processed with the Fiji stitching plugin (http://www.discotechnologies.org/).

### Reconstructions of full-body scans and quantification

Detailed step-by-step instructions for image data stitching and volume fusion were provided previously6. Briefly, image stacks were recorded using ImSpector software (LaVision BioTec GmbH) and saved in TIFF format for each channel separately. The scanned ventral and dorsal-mouse image data were first stitched using the Fiji stitching plugin and volumes fused using Vision4D (v3.5 x64, Arivis AG, version 3.4.0). To increase the precision of volume fusion, alignment was performed by manually selecting 3 to 4 anatomical landmarks from the overlapping regions. Representative images were created using Imaris (Bitplane AG, version 9.6.0) and Vision4D for 3D volumetric reconstruction, maximum intensity projection, and depth color rendering. To isolate a specific tissue region, the Imaris surface tool was used manually and the mask channel option for pseudocolor was selected. After manual segmentation, the region was visualized in 2D slices using the Ortho Slicer tool. Virtual reality (VR) pictures and movies were generated using the Syglass software (IstoVisio, Inc, version 1.7.2).

For quantification of the myenteric plexus in the duodenum between germ free mice and wild type mice, five 200 μm × 200 μm × 200 μm cubic volumes along the portal triads were randomly selected from the reconstructed 3D images in Imaris. The length of the PGP 9.5-positive myenteric plexus in each cubic volume was traced using Imaris Filament Tracer.

### Virtual Reality (VR) Headset procedure

Movies 24-33 require a virtual reality (VR) headset. To visualize them, you need a VR-video player on your VR-device or computer. Videos played in virtual reality need to have “_360” at the end of their file name and set to a “360°/3D” view in the VR-player for an immersive experience.

### Quantification

Data are presented as mean ± s.d. Statistical analysis was performed using Prism GraphPad software v.6 with 95% confidence interval. P values were calculated using two-tailed unpaired t test to compare data between two groups. P values of <0.05 were considered statistically significant.

## ACKNOWLEDGEMENTS

Illustrations were created with BioRender.com. H.M. would like to thank the China Scholarship Council (CSC) for the financial support (No. 201806780034). We thank Dr. Lun Peng (department of pharmacy, LMU Munich) and Dr. Yi Lin (department of pharmacy, LMU Munich) for dynamic light scattering measurements. We thank Dr. Markus Elsner for editing the manuscript.

This work was supported by the Vascular Dementia Research Foundation, Deutsche Forschungsgemeinschaft (DFG, German Research Foundation) under Germany’s Excellence Strategy within the framework of the Munich Cluster for Systems Neurology (EXC 2145 SyNergy, ID 390857198) and DFG (SFB 1052, project A9; TR 296 project 03) as well as German Federal Ministry of Education and Research (Bundesministerium für Bildung und Forschung, BMBF) within the NATON collaboration (01KX2121).

## AUTHOR CONTRIBUTION

A.E., conceived and led all aspects of the project. H.M., and J.L., developed the method and conducted most of the experiments. R.A.M., I.H., J.C.P., and M.T., analyzed data. L.H. generated VR videos. A.E., and F.H., supervised the project. J.L. and H.M., wrote the first draft of manuscript. A.E. wrote the final manuscript. All authors reviewed and approved the final manuscript.

## MOVIE LEGENDS

**Movie S1**. 3D reconstruction of a mouse labeled with PGP 9.5 and imaged with light-sheet microscopy. Different PGP 9.5 innervation regions (green) are visualized with high contrast over background (gray).

**Movie S2**. 3D view of PGP 9.5 positive peripheral nerve in mouse heart showing in magenta.

**Movie S3**. 3D visualization of sympathetic nerve in mouse spleen labeled with tyrosine hydroxylase (TH) in magenta.

**Movie S4**. 3D visualization of mouse liver and gallbladder innervated with PGP 9.5 positive peripheral nerves in magenta.

**Movie S5**. 3D annotation of mouse intestinal innervation with TH positive nerve in magenta. Grid-like lattice structures can be seen in the intestinal wall.

**Movie S6**. 3D annotation of neuronal connections in multiple organs (kidney, spleen, liver, and intestine) labeled with PGP 9.5 in green.

**Movie S7**. 3D visualization of the entire lymphatic vessels in the whole mouse. The lymphatic vessels of a 4-week-old mouse were labeled with LYVE1 in yellow.

**Movie S8**. 3D visualization of the hind limb of a mouse with LYVE1 lymphatic vessel labeling in yellow. Fine details of the lymphatic vessels and lymph nodes are shown.

**Movie S9**. 3D visualization of mouse kidney with LYVE1 lymphatic vessels labeled in yellow.

**Movie S10**. 3D annotation of mouse stomach with LYVE1 lymphatic vessels highlighted in yellow. The fine details of the lymphatic vessels are clearly visible throughout the scan.

**Movie S11**. 3D annotation of mouse intestine lymphatic vessels with yellow color labeled by LYVE1 and scanned with light-sheet microscopy. Fine details of the lymphatic vessels can be seen in the intestine and a lymph node is located adjacent to the intestine.

**Movie S12**. 3D view of the intestinal wall of a mouse with LYVE1 lymphatic vessels labeled in yellow.

**Movie S13**. wildDISCO staining of PROX1 lymphatic vessels in green and arterial staining of alpha-SMA in red, revealing lymphatic vessels penetrated in mouse cerebral cortex.

**Movie S14**. 3D illustration showing wildDISCO staining of LYVE1 in yellow and podoplanin in magenta to visualize lymphatic capillaries covered on the surface of the mouse brain, and also entering the brain parenchyma around thalamus.

**Movie S15**. 3D illustration showing innervation of intestinal lymph nodes by sympathetic neurons using TH staining to label sympathetic nerves in magenta and CD45 staining to label immune cells in green.

**Movie S16**. Representative 3D reconstructions of immune cells on the intestine neurons. CD45 is stained in green for immune cell distribution and TH stained in magenta for sympathetic nerves.

**Movie S17**. 3D reconstruction of a TH and LYVE1 labeled mouse by light sheet microscopy. Different regions of sympathetic nerve innervation in green and lymphatic vessels in blue can be seen.

**Movie S18**. 3D illustration of the TH stained sympathetic nerve in green interacting with LYVE1-labeled lymphatic vessels in yellow on the intestinal wall.

**Movie S19**. 3D illustration of TH+ sympathetic nerve (green) interacting with LYVE1+ lymphatic vessels (yellow) inside the intestine.

**Movie S20**. 3D illustration showing the distribution of multiple lymph nodes throughout the mouse. Innervation of pan-neuron markers PGP 9.5 in green and lymph node masked color in cyan.

**Movies S21**. Representative 3D illustration of a posterior limb lymph node innervated by PGP 9.5 positive nerves in green and PROX1 positive lymph nodes in magenta.

**Movie S22**. 3D illustration showing PGP 9.5 staining of myenteric nerves in green from germ-free mouse gut. In germ-free mice, the myenteric plexus appears disorganized in some regions of the gut wall, with fewer ganglia and thinner nerve connecting fibers.

**Movie S23**. Summary of results showing a homogeneous and simultaneous antibody staining throughout the entire mouse bodies.

**Movie S24**. VR 3D visualization of neuronal connections in multiple organs labeled with PGP 9.5 in green and masked colors for kidney, spleen, liver, and intestine.

**Movie S25**. VR 3D visualization of mouse intestinal innervation with TH positive nerve in magenta referred to Movie S5. Grid-like structures can be clearly visible throughout the intestine.

**Figure S1.**
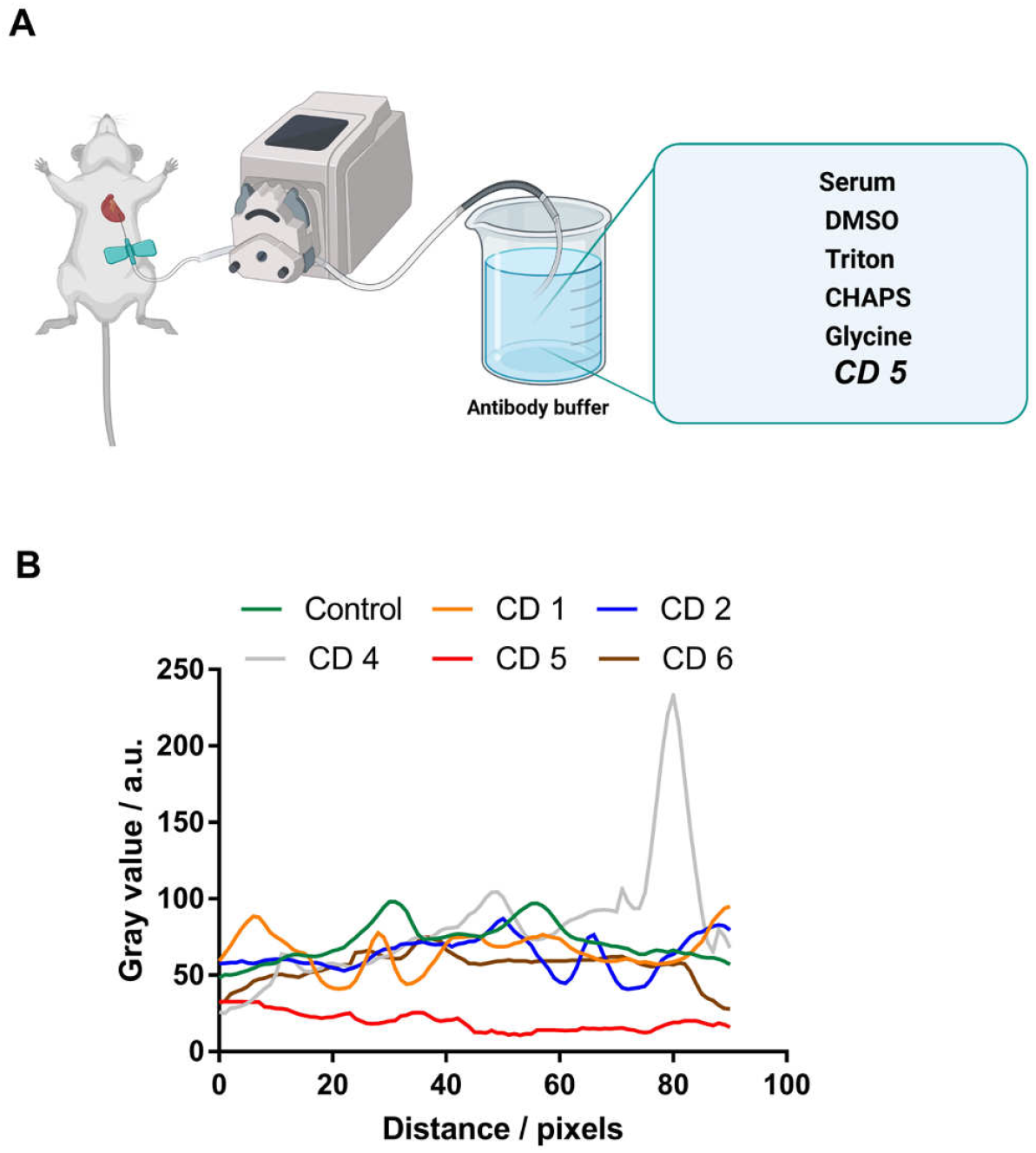
Overview of wildDISCO immunostaining buffer and quantification of permeabi-lization efficacy. (A) Illustration of primary chemical components involved in the wildDISCO immunostaining buffer. (B) Profile plot along each mouse brain dimension in Fig. 1C.

**Figure S2.**
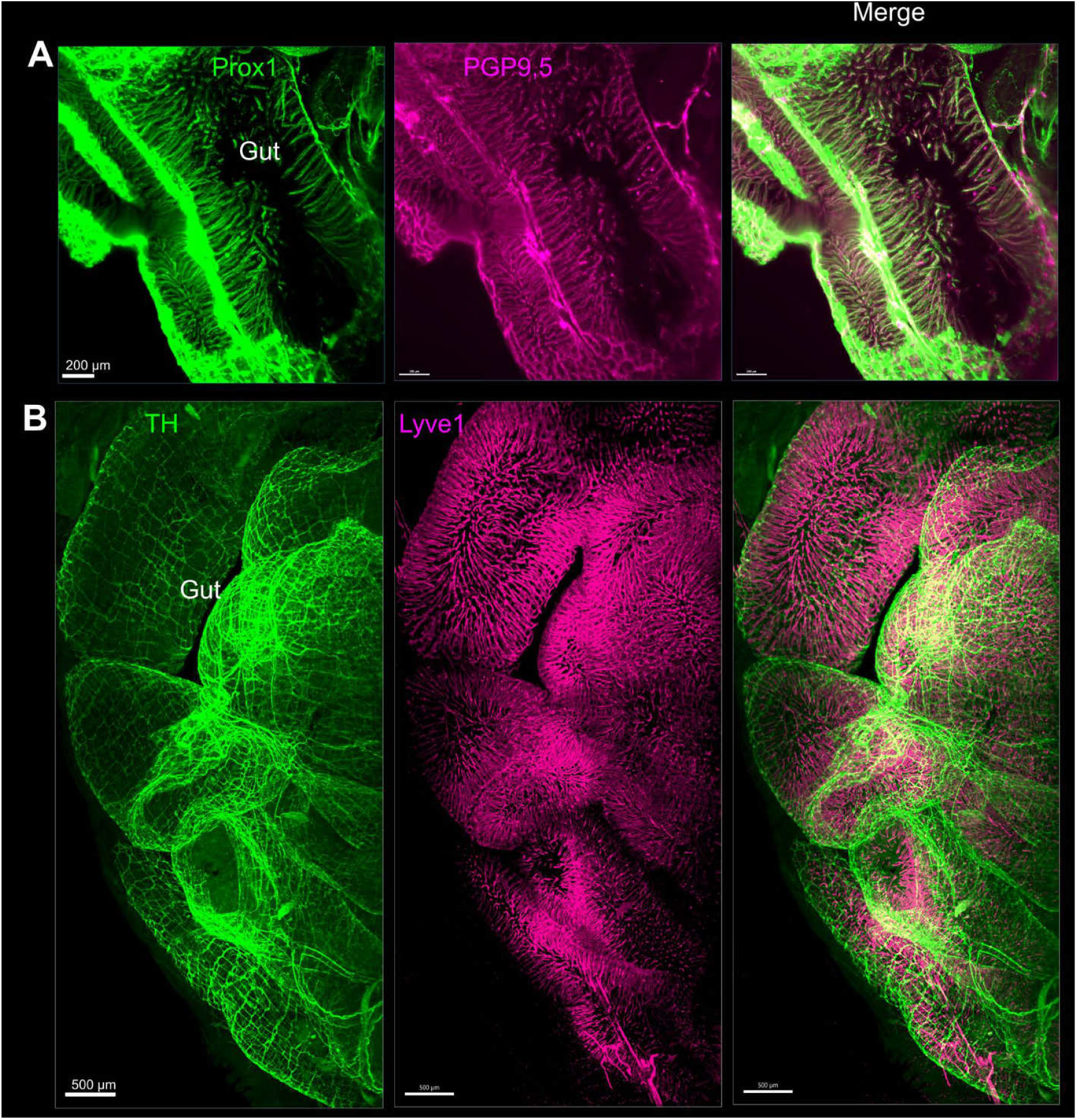
Nerve-lymphatic vessel interactions in the gut. (A) PGP9.5 nerve fibers (magenta) interacted with Prox1 lymphatic vessel (green). (B) TH sympathetic nerve (green) and LYVE1 lymphatic vessel (magenta).

**Figure S3.**
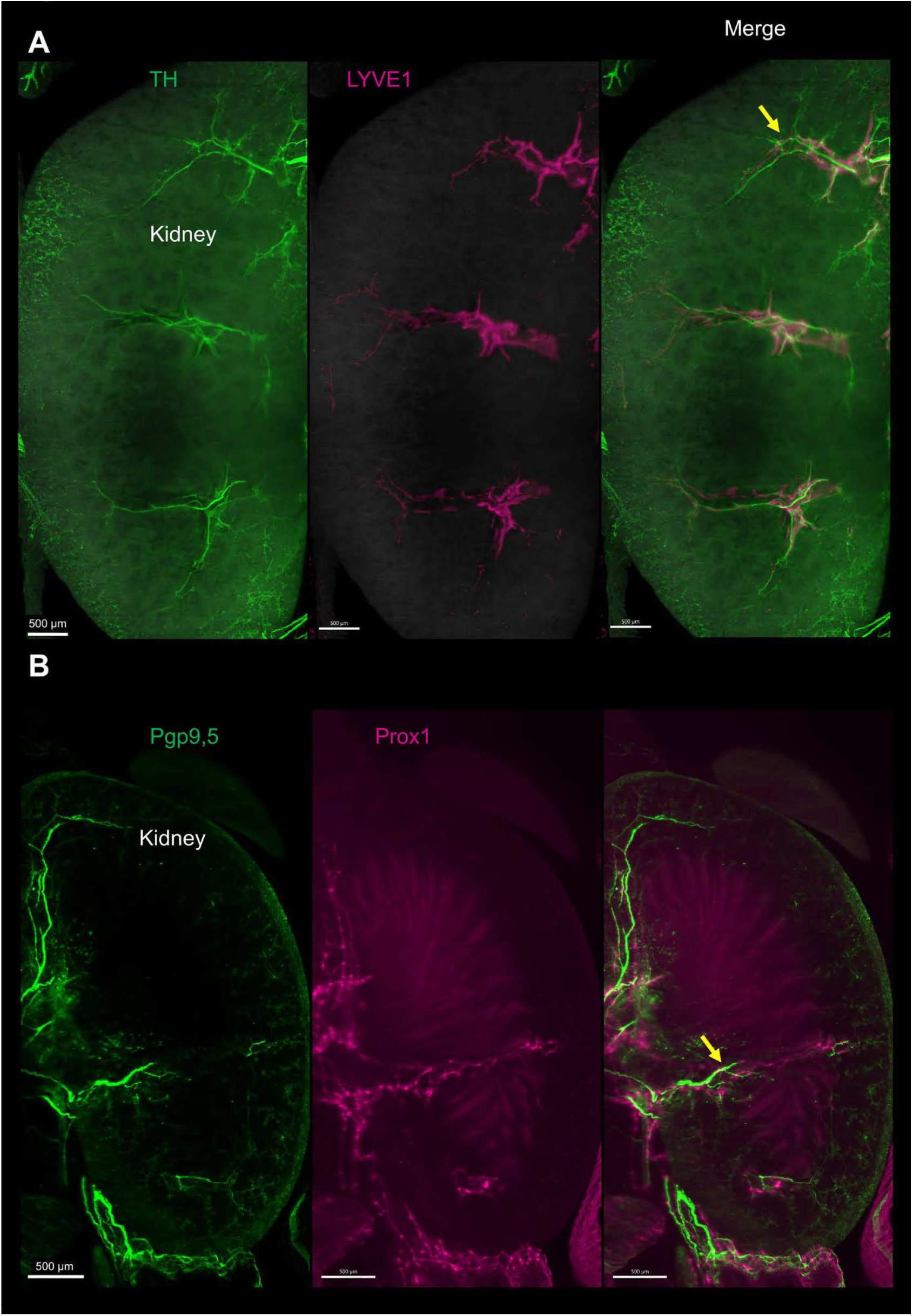
Nerve-lymphatic vessel interactions in the kidney. (A) The TH sympathetic neuron (green) innervated the LYVE1 lymphatic vessel (magenta) in the kidney. (B) The PGP 9.5 pan-neuronal marker (green) combined with Prox1 lymphatic vessel marker (magenta).

